# Host defense mechanisms induce genome instability in an opportunistic fungal pathogen

**DOI:** 10.1101/2021.04.01.438143

**Authors:** Amanda Smith, Meleah A. Hickman

## Abstract

The ability to generate genetic variation facilitates rapid adaptation in stressful environments. The opportunistic fungal pathogen *Candida albicans* frequently undergoes large-scale genomic changes, including aneuploidy and loss-of heterozygosity (LOH), following exposure to physiological stressors and host environments. However, the specific host factors that induce *C. albicans* genome instability remains largely unknown. Here, we leveraged genetically-tractable nematode hosts to specifically investigate the innate immune components driving host-associated *C. albicans* genome instability, which include host production of antimicrobial peptides (AMPs) and reactive oxygen species (ROS). *C. albicans* associated with wildtype, immunocompetent hosts carried multiple large-scale genomic changes including LOH, whole chromosome, and segmental aneuploidies. In contrast, *C. albicans* associated with immunocompromised hosts deficient in AMPs or ROS production had reduced LOH frequencies and fewer, if any, additional genomic changes. We also found that *C. albicans cap1*Δ/Δ strains deficient in ROS detoxification, were more susceptible to host-produced ROS genome instability compared to wildtype *C. albicans*. Further, genomic perturbations resulting from host-produced ROS are mitigated by the addition of antioxidants. Taken together, this work suggests that host-produced ROS and AMPs induces genotypic plasticity in *C. albicans* which may facilitate rapid adaptation and lead to phenotypic changes.

## Introduction

The opportunistic fungal pathogen, *Candida albicans*, is a leading cause in fungal bloodstream infections, and 40% of these infections result in mortality^1^. In addition to bloodstream infections, *C. albicans* causes non-lethal mucosal infections, including vaginal and oral candidiasis^1^. Despite its ability to cause infection, *C. albicans* is typically commensal and a component of the human microbiome^2^. The ability of *C. albicans* to cause infection is highly dependent on the host context, including high estrogen levels^3^, chronic stress^4^, antibiotic use^5–7^, uncontrolled diabetes^8, 9^, and immunosuppression^10, 11^. In the absence of proper immune recognition, fungal proliferation is uncontrolled, leading to infection. However, despite having a fully functioning immune system, healthy individuals also experience *C. albicans* infections. This is often due to commensal isolates within an individual that become pathogenic rather than an infection from outside sources^12^. This transition may be facilitated by *C. albicans* phenotypic and genotypic heterogeneity. *C. albicans* genetic diversity within and among individuals is very high and often include numerous single nucleotide polymorphisms (SNPs) and loss of heterozygosity (LOH) events ^13^. This genetic variation may be a direct consequence of the stressors *C. albicans* encounters in the host which include immune stressors and other microbes. Recent work has demonstrated the host environment elevates genome instability in *C. albicans* similar to *in vitro* stressors ^14–18^. However, the specific host components that generate this instability is largely unknown.

Several studies investigating the diversity of clinical *C. albicans* isolates found that genomic variation was quite common^13, 19^. This variation included polymorphisms, copy number variation, loss-of-heterozygosity (LOH), and partial or whole chromosomal aneuploidies^19^. This suggests that the host environment rapidly generates genetic variation. Recent studies involving murine infection models have found that when exposed to different host niches, *C. albicans* has higher levels of genetic variation compared to *in vitro*^14–17^. Similar to clinical isolates, laboratory *C. albicans* often had large-scale genomic changes including aneuploidy and LOH following murine infection^14–16^. In addition to murine models, *Caenorhabditis elegans* has been used as a model to assess *C. albicans* genome stability and strains that differ in genetic background and ploidy have elevated host-associated genome instability^17^. It is clear that host environments drive genetic diversity in *C. albicans,* however, it is remains unclear what specific host components contribute to *C. albicans* genome instability.

In the host, *C. albicans* likely encounters many different stressors within the host environment, including the immune system which is designed to control and remove pathogens. The human immune system recognizes pathogens, like *C. albicans* through pathogen recognition receptors (PRRs) that detect the specific microbial chemical signatures called pathogen-associated molecular patterns (PAMPS)^20^. The main cells involved in the recognition of *C. albicans* include monocytes, macrophages, and neutrophils^21^. This recognition triggers a specific cellular response. For instance, *β*-glucan is primarily recognized through CR3 and Dectin-1 which induces the production of antimicrobial peptides (AMPs) and pro-inflammatory cytokines^22^. AMPs inhibit microbial growth through a variety of methods, including disruption of the cell membrane, and halt DNA, RNA, and protein synthesis^23^. The release of AMPs and pro-inflammatory cytokines recruit other immune cells to the site of infection including neutrophils^21, 22^. Upon phagocytosis of *C. albicans*, neutrophils and macrophages produce an oxidative burst, which generates reactive oxygen species (ROS)^21, 22^. ROS cause cellular toxicity through structural changes to the DNA^24^ and in *C. albicans* generates double strand breaks (DSBs)^25^. Together, host-produced AMPs and ROS act in various ways, damaging the cell membrane, and causing DNA damage in order to kill *C. albicans* cells.

*C. albicans* has several mechanisms to mitigate damage from host-produced ROS and AMPs, including secretion of peptide effectors^26^, induction of the HOG1 stress response pathway, the RAD53 DNA damage checkpoint, and upregulation of the transcription factor cap1p^27^. Cap1p activates antioxidant genes that break down ROS^27^. Despite these mechanisms, *C. albicans* has a higher frequency of loss-of-heterozygosity^28^ following exposure to hydrogen peroxide *in vitro*. Similarly, tetraploid *C. albicans* acquire DSBs and undergo ploidy reduction in response to ROS^25^. However, we still don’t understand the impact of host-produced ROS or AMPs on *C. albicans* genome stability.

To specifically investigate the impact of host-produced ROS and AMPs on *C. albicans* genome stability we can utilize the nematode, *Caenorhabditis elegans*. *C. elegans* is an extremely powerful model for investigating *C. albicans* pathogenesis and genome stability. In addition to its rapid life cycle, easy maintenance, and large brood sizes, it has a highly conserved innate immune system which includes both AMPs and ROS^29^. In response to *C. albicans* infection, *C. elegans* produce a majority of AMPs through the mitogen-activated protein kinase (MAPK) signaling cascade^30, 31^. This pathway includes SEK-1 (MAPKK), homologous to the MKK3/6 and MKK4 mammalian MAPKKs. Mutations in SEK-1 increase susceptibility to *C. albicans* as well as other microbial pathogens demonstrating its importance in defense^32, 33^. In addition to AMPs, *C. elegans* produce ROS as another host defense, similar to human macrophages. ROS are produced in response to bacterial and fungal pathogens and is mediated by the dual oxidase BLI-3^34–37^. However, when BLI-3 is mutated, *C. elegans* are more susceptible to *C. albicans*^38^.

Here we investigated the role host-produced AMPs and ROS play in generating genome instability in *C. albicans* using the model host *C. elegans*. We infected healthy and two different immunocompromised hosts with mutations in either *sek-1* (AMP production) or *bli-3* (ROS production) with *C. albicans* and subsequently measured LOH frequency and genome-wide changes with whole-genome sequencing. We found that healthy hosts elevate genome instability and generate greater genetic diversity in *C. albicans* compared to immunocompromised hosts. Furthermore, our results suggest that the dual oxidase, BLI-3 produces genome instability through the production of ROS. The genome instability caused by host-produced ROS can however be alleviated with the addition of antioxidants. Taken together, our results suggest that host innate immune pathways cause genome instability in *C. albicans.* The genome diversity generated inside healthy hosts likely provide *C. albicans* with the ability to quickly adapt to a changing and stressful host environment. Further, these genome-wide perturbations may be a way in which *C. albicans* transitions from commensal to pathogenic in immunocompetent hosts.

## Results

### Host defense pathways elevate C. albicans genome instability

We, and others, have shown that nematode and murine host environments induce *C. albicans* genome instability compared to *in vitro*^14–18^, but the specific host attributes driving genome instability have yet to be elucidated. Here, we tested whether components of host innate immune function contributed to host-associated genome instability by comparing the frequency of loss-of-heterozygosity (LOH) between *C. albicans* extracted from healthy and immunocompromised hosts (Fig. 1A). We used two different immunocompromised host genotypes, one carrying a deletion of *sek-1* which cannot produce AMPs^39^, and another host genotype carrying a deletion of *bli*-3 which cannot produce ROS^35, 37^. Host-associated LOH frequencies were reduced in *C. albicans* extracted from both *sek-1* and *bli-3* hosts compared to *C. albicans* extracted from healthy hosts (Fig. 1B). Following host exposure, the relative frequency of LOH of both laboratory and clinical *C. albicans* was significantly lower in *sek-1* and *bli-3* hosts compared to wildtype (N2) hosts (Figs. 1B, S1A&B). Our previous work demonstrated that the bloodstream strain was more unstable compared to the laboratory strain. Despite this higher baseline instability, we found a significantly lower LOH frequency of bloodstream *C. albicans* extracted from wildtype hosts compared to *bli-3* hosts (Fig. S1B).

**Figure 1:**
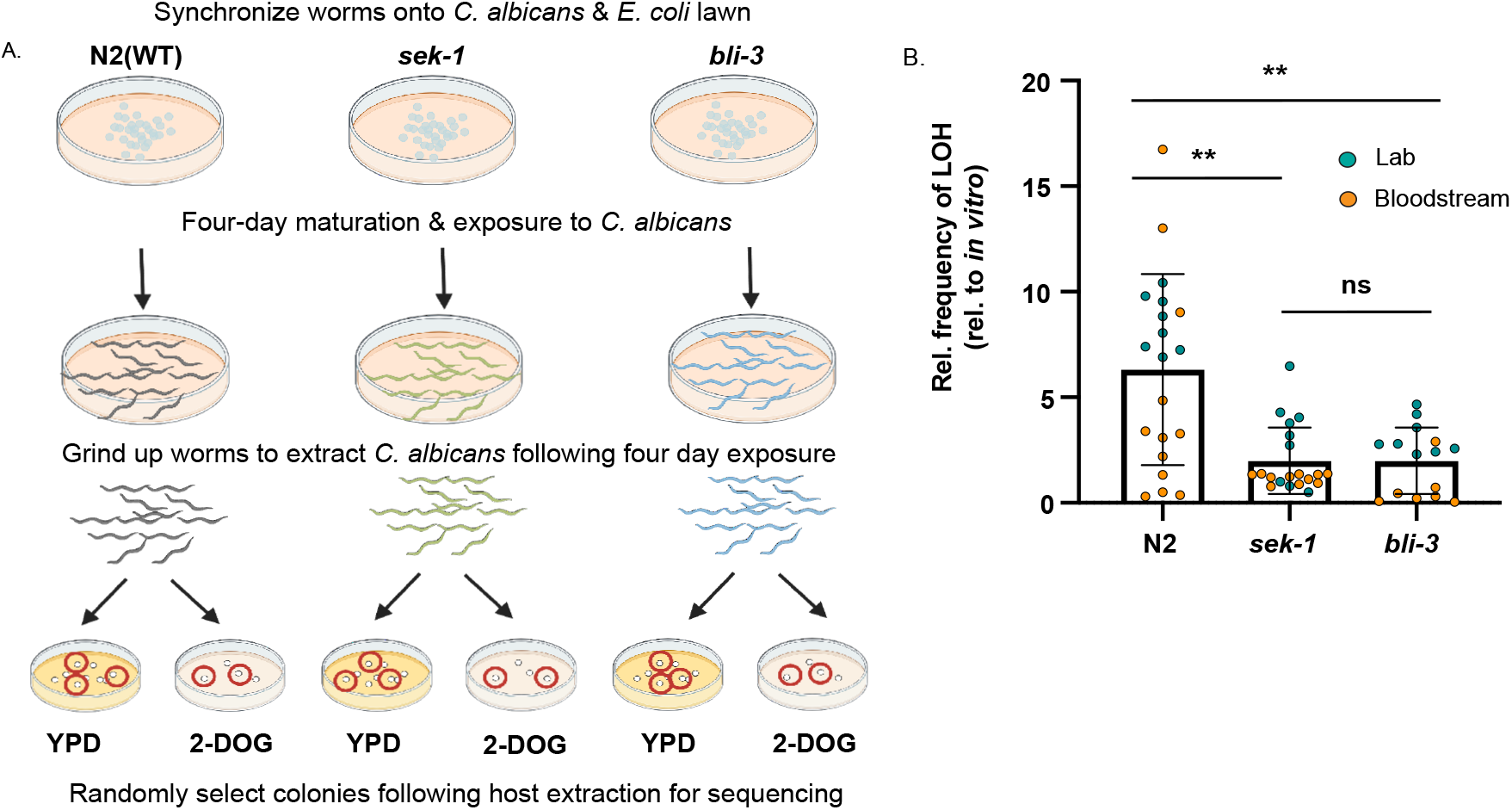
Host immunity impacts *C. albicans* genome stability. **A)** Experimental overview of in vivo experiments. **B)** Laboratory (teal) and bloodstream (orange) *C. albicans* LOH frequency following host association relative to the LOH frequency of *C. albicans* no host control for N2, *sek-1*, and *bli-3*. Plotted are the means and SD. Asterisks indicate significant differences (**** p < 0.0001, ns = not significant; Kruskal-Wallis with post-hoc Dunn’s multiple test).

Together, these data suggest that *bli-3* and *sek-1* immune pathways may be a source of genome instability in *C. albicans* regardless of pathogen genetic background.

### Host immunity impacts the genomic landscape of C. albicans

While LOH assays are an easy and useful way to measure genome instability in *C. albicans,* they only provide a narrow focus into one small part of the genome. Therefore, we wanted to look more broadly and determine how host immunity impacts the genomic landscape of *C. albicans*. Following host-association and extraction from wildtype, *sek-1* and *bli-3* hosts we randomly selected single colonies from both YPD (no LOH selection) and 2-DOG (LOH, selection) and performed whole-genome sequencing. Large-tract LOH events and whole-chromosomal and segmental aneuploidy were frequently observed in LOH-selected isolates from wildtype hosts, but not from either immunocompromised host background (Fig. 2). Whole or segmental aneuploidy was observed in 66% (4/6) the LOH-selected isolates following wildtype host-association, but never from immunocompromised hosts. One isolate was trisomic for both chromosome 6 and 7. Of the additional LOH tracts not selected for, all were caused by break-induced recombination. In contrast, we did not detect any aneuploidy in the isolates from either of the immunocompromised hosts (Fig. 2). We also investigated the impact of host exposure on the genomic landscape of isolates with no prior selection and found 50% (2/4) isolates had LOH events during exposure to healthy hosts (Fig. S2A). However, we did not detect any LOH events in the isolates extracted from immunocompromised hosts (Fig. S2A). Thus, healthy hosts induce more large-scale genomic changes in *C. albicans* regardless of LOH selection, compared to immunocompromised hosts.

**Figure 2:**
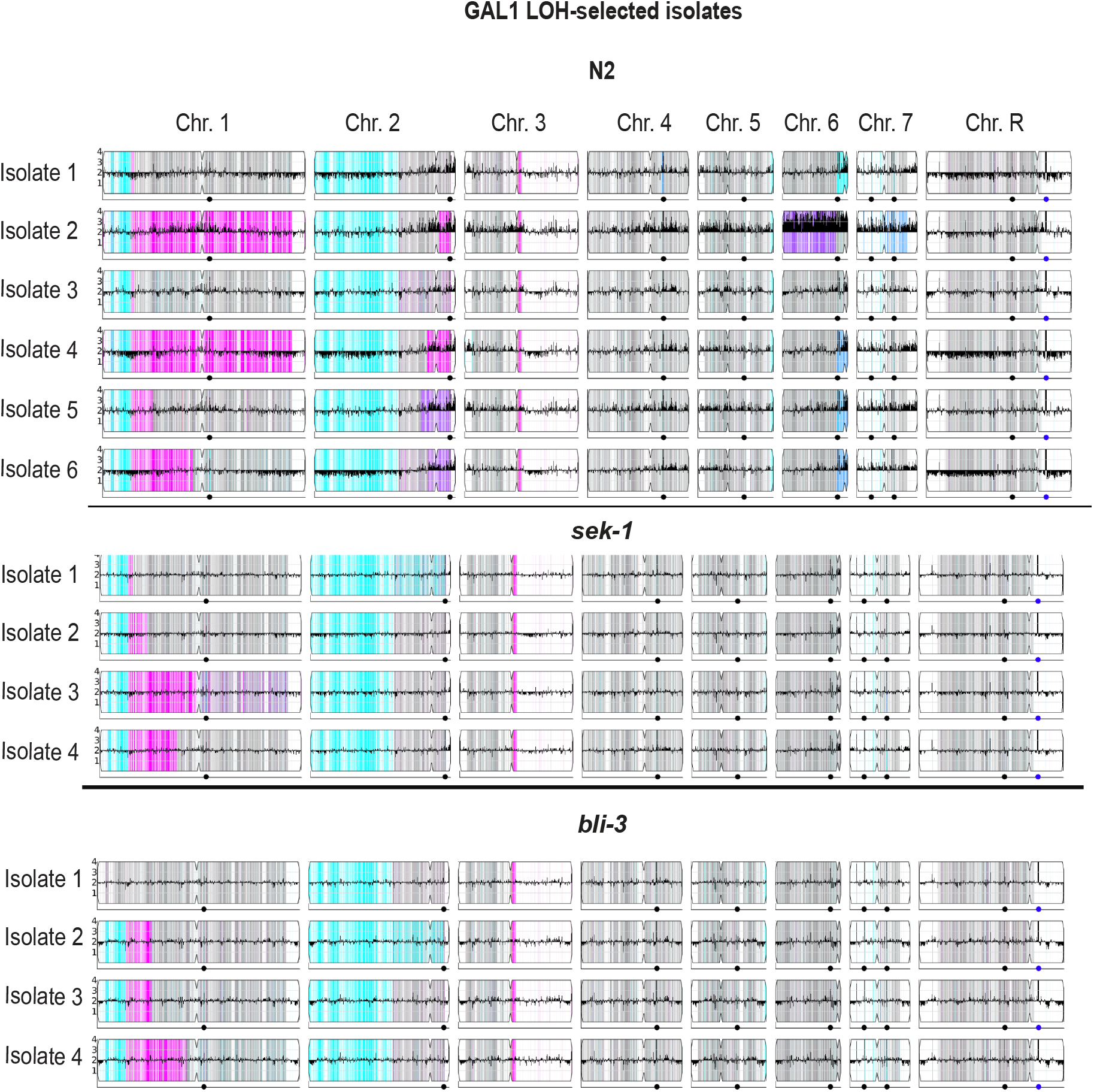
Genome-wide changes following host association. YMAPs showing the copy number and allelic ratio for each GAL-isolate following host exposure.

We also investigated whether exposure to wildtype hosts generate more small-scale genetic changes in *C. albicans* compared to immunocompromised hosts. PCA analysis shows very little variation between the isolates, regardless of LOH selection or host background (Fig. S2B), thus we grouped LOH selected and non-LOH selected isolates together for each host background and compared the number of SNPs and INDELS. Because ROS are mutagenic, we expected more mutations in *C. albicans* extracted from hosts and ROS. However, we detected no significant difference in the number of SNPs in the isolates extracted from the healthy or immunocompromised hosts (Fig. 3A). We did detect a significantly lower number of INDELs (~1000 less) in isolates extracted from *bli-3* hosts compared to *sek-1* hosts (Fig. 3B). This suggests that the BLI-3 defense pathway generates INDELs in *C. albicans*. Despite incurring roughly the same number of mutations regardless of host background, larger chromosomal events including aneuploidy and large-scale LOH are primarily observed in isolates from wildtype hosts and not immunocompromised hosts.

**Figure 3:**
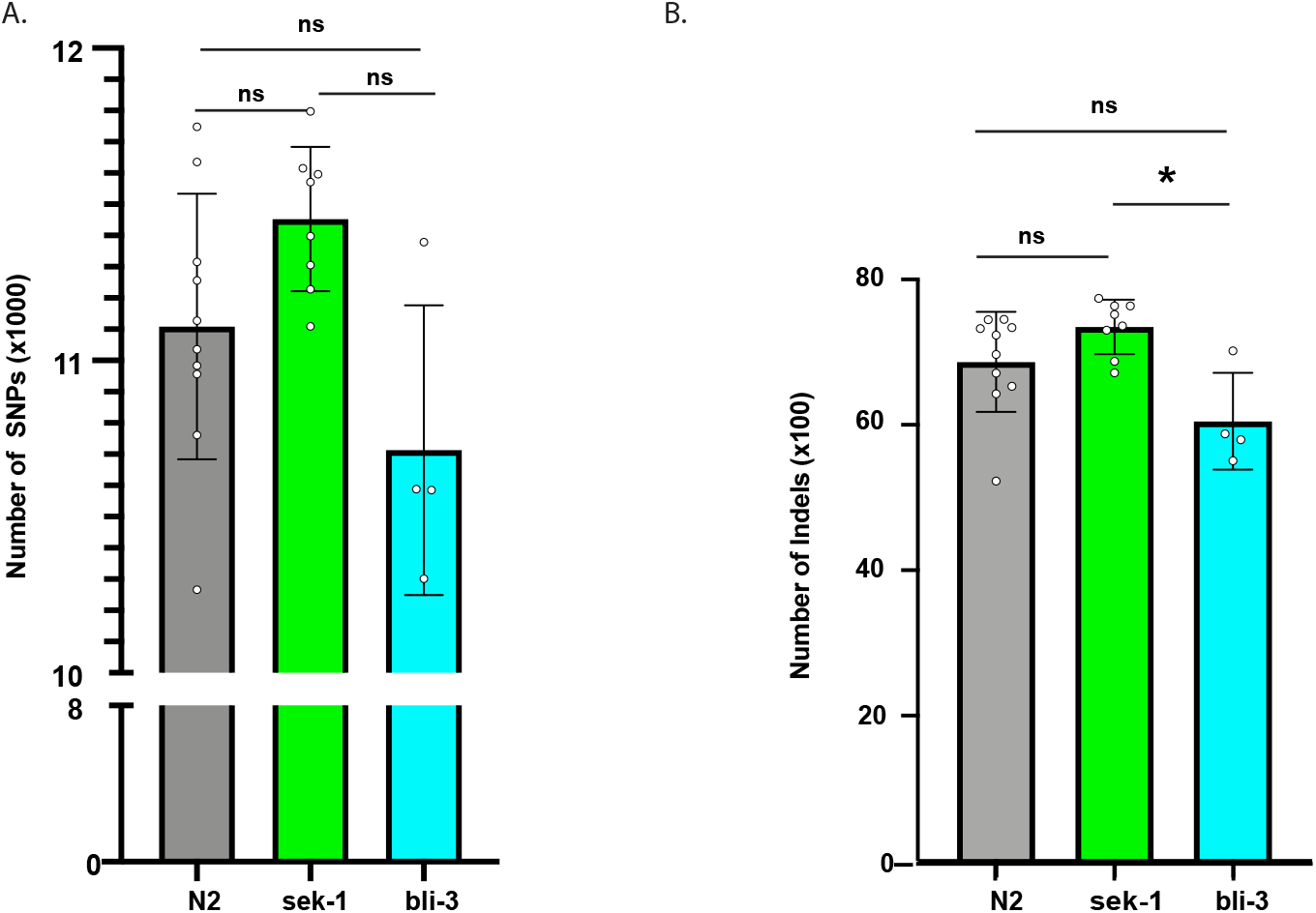
*bli-3* hosts generate less genome-wide INDELs in *C. albicans.* **A)** Number of genome-wide SNPs (x1000) in *C. albicans* following exposure to N2 (wildtype, grey, n = 10), *sek-1* (green, n = 8), and *bli-3* (blue, n = 4) hosts. Plotted are the mean and SD. Individual data points represent the number of SNPs in one isolate. **B)** Number of genome-wide INDELs (x100) in *C. albicans* following exposure to N2 (wildtype, grey, n = 10), *sek-1* (green, n = 8), and *bli-3* (blue, n = 4) hosts. Plotted are the mean and SD. Individual data points represent the number of INDELs in one isolate. Asterisks indicate significant differences (* p < 0.05, ns = not significant; ANOVA with post-hoc Tukey’s multiple test).

Our data suggest that both *sek-1* and *bli-3* host immune pathways are a source of pathogen genome instability. While the SEK-1 MAPKK plays a central role in the MAPK cascade with many downstream targets, the main role of BLI-3 is generating ROS. Because of the clear connection between BLI-3 and ROS production, we wanted to investigate whether host-produced ROS is generating genome instability in *C. albicans*. We exposed both wildtype (SC5314) *C. albicans* and a *cap1Δ/Δ* mutant strain of *C. albicans* deficient in detoxifying ROS, to 5 mM H_2_O_2_ for 24 hours *in vitro* and measured the frequency of LOH. Similar to previous literature 28, we found exposure to *in vitro* H_2_O_2_ increased LOH frequencies in both wildtype and *cap1Δ/Δ C. albicans* strains by 15- and 40-fold, respectively, when compared to the no stress treatment (Fig. 4A & S3). We found that the relative LOH frequency following exposure to H_2_O_2_ was significantly higher in *cap1Δ/Δ C. albicans* compared to wildtype *C. albicans.* To directly test if host-produced ROS elevates *C. albicans* genome instability, we compared *cap1Δ/Δ C. albicans* LOH frequency to wildtype *C. albicans* LOH frequency following host exposure. If host-produced ROS elevate *C. albicans* LOH frequency, we expect *cap1Δ/Δ C. albicans* to have a higher frequency of LOH after exposure to all hosts except for *bli-3* hosts, which do not produce ROS. The *cap1Δ/Δ* strain had significantly higher host-associated LOH frequencies from all host backgrounds compared to *in vitro* (Fig. S3B). The *cap1Δ/Δ* strain had significantly higher relative LOH frequency compared to wildtype *C. albicans* extracted from both healthy and *sek-1* hosts (Fig. 4B). Unsurprisingly, we did not detect any significant differences in the relative LOH frequency between wildtype *C. albicans* and *cap1Δ/Δ C. albicans* extracted from *bli-3* hosts which do not produce ROS(Fig. 4B). Together, these data indicate that host-produced ROS elevates *C. albicans* genome instability and cap1p is important in detoxifying both *in vivo* and *in vitro* ROS.

**Figure 4:**
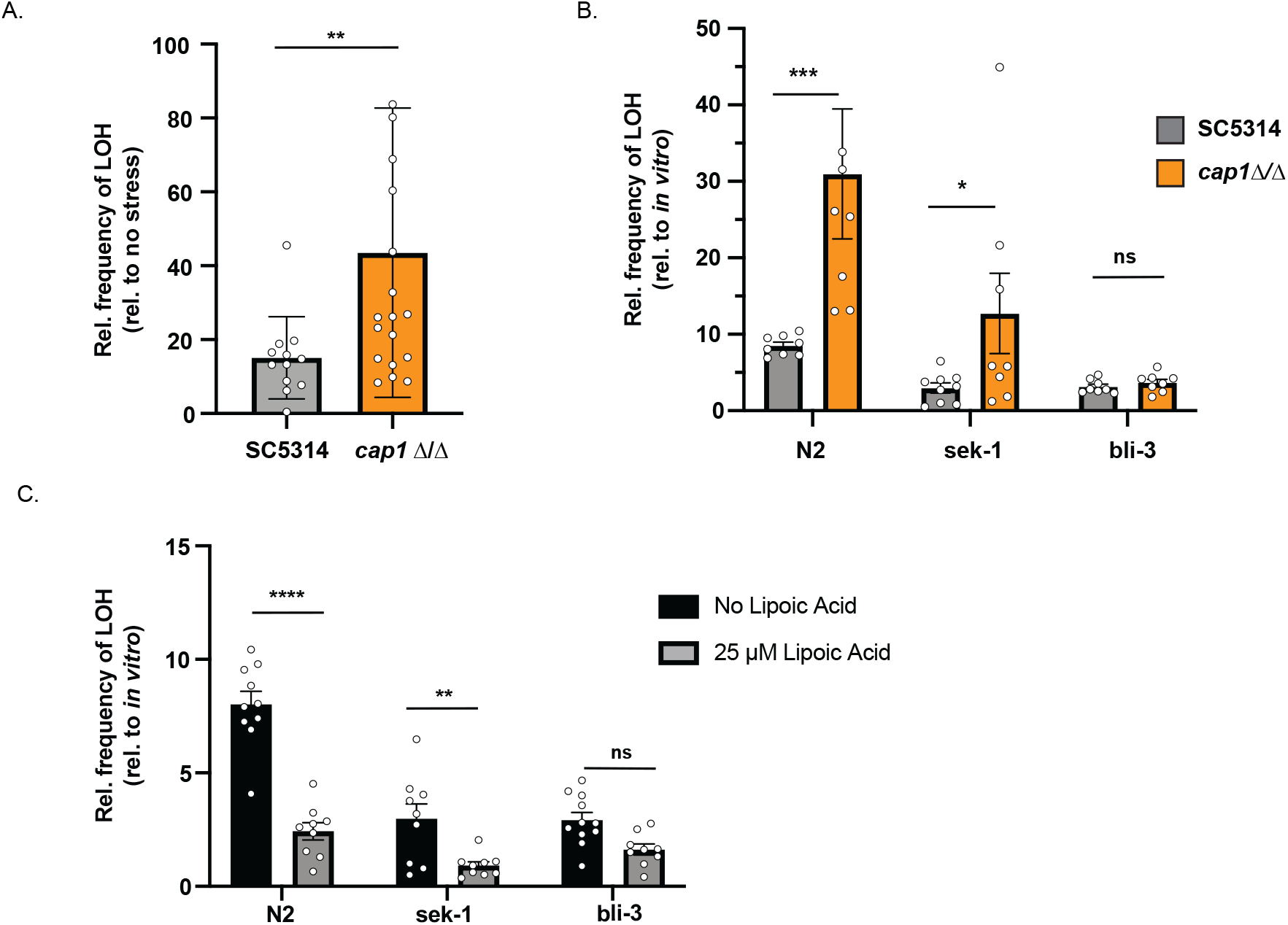
*C. albicans cap1* Δ/Δ strain is more susceptible to *in vitro* and *in vivo* ROS. **A)** Plotted are the LOH frequencies of *C. albicans* exposed to 5mM H_2_O_2_ for 24 h relative to the frequencies of LOH of *C. albicans* without stress exposure. Plotted are the means and SD for both SC5314 (grey, n = 12) and *cap1*Δ/Δ (orange, n = 19). Each data point represents an individual measurement. **B)** Plotted are the LOH frequencies of *C. albicans* exposed to each host environment relative to the no host frequency of LOH. Plotted are the means and SD. **C)** *C. albicans* LOH frequency following host exposure relative to the no host LOH frequency with 25 μM of alpha-lipoic acid (grey, N2: n = 9, *sek-1*: n = 9, *bli-3*: n = 9) and without 25 μM of alpha-lipoic acid (black, N2: n = 16, *sek-1*: n = 9, *bli-3*: n = 19). Each data point represents an individual measurement. Plotted are the means and SD. Asterisks indicate significant differences (**** p < 0.0001, *** p < 0.005, ** p < 0.01, * p < 0.05, ns = not significant; Mann-Whitney U test).

Since ROS generated through the dual oxidase, BLI-3 causes genome instability in *C. albicans,* we wanted to determine whether antioxidants mitigate *C. albicans* genome instability from host-produced ROS. To test whether antioxidants reduce *C. albicans* genome instability caused by host-produced ROS, we measured *C. albicans* LOH frequency following host exposure with the addition of 25 μM of lipoic acid ^37^, a well research antioxidant involved in the breakdown of ROS. In *C. albicans* extracted from healthy hosts, we observed a significant reduction in relative LOH frequency with the addition of lipoic acid (Figs. 4C & S5C). We observed a similar trend in *C. albicans* extracted from *sek-1* hosts. Lipoic acid lowered the relative frequency of LOH in *sek-1* hosts (Figs. 4C & S5C). Following *bli-3* host exposure *C. albicans* relative LOH frequency was not significantly different with or without lipoic acid (Figs. 4C & S5C). This suggests that antioxidants only reduce *C. albicans* genome instability in hosts that produce ROS. Together, these data suggest that the host-produced ROS generates genome instability in *C. albicans* which can be alleviated by the use of antioxidants.

## Discussion

We previously reported higher levels of genome instability in *C. albicans* exposed to wildtype hosts^17^. Here we investigated whether host innate immune pathways impact *C. albicans* genome stability. By using wildtype and two immunocompromised hosts with mutations in AMP production (*sek-1*) and ROS production (*bli-3*), we compared the differences in host-associated *C. albicans* genome stability and mutational landscape. *C. albicans* associated with either immunocompromised host had reduced relative LOH frequencies compared to those associated with healthy hosts (Figure 1). Similar to several other host passaging experiments and whole-genome sequencing results from clinical isolates^14, 15, 19^, many of our isolates (5/6) extracted from wildtype hosts contained large-scale genomic changes including whole and segmental chromosomal aneuploidy and/or additional LOH events (Figure 2). Akin to Forche et. al.,^14^ we detected the presence of an extra copy of chromosome 6 following host exposure. Their study suggests that this extra copy of chromosome 6 produces a more commensal-like phenotype inside the host environment. However, we have yet to investigate the virulence of our isolates. We also detected an extra copy of chromosome 7 in one isolate following wildtype host exposure, which in a gastrointestinal murine model of candidiasis is more fit in the gastrointestinal tract compared to the euploid parent^15^. Therefore, the generation of specific aneuploidies might enable host adaptation. However, removal of host-produced AMPs and ROS lead to the overall decrease in genetic variation. Immunocompromised host-associated *C. albicans* isolates did not carry any detectable aneuploidy and only a small number (2/8) had an LOH event that was not selected for (Figure 2). This suggests that both AMPs and ROS act as stressors on *C. albicans* that enable the generation of genetic variation which might lead to phenotypic changes that create a more synergistic host–pathogen relationship.

The host has a variety of mechanisms in order to control microbial growth. Because *C. albicans* is normally a commensal and resides in many niches in the human body, it must strike a delicate balance with the host to evade detection. Overgrowth of *C. albicans* initiates an immune response that initially includes the production of AMPs and the recruitment of phagocytes to the site of infection^21^. AMPs represent a potent defense mechanism designed to kill pathogens through a variety of mechanisms including disruption of the cell membrane and DNA damage^23^. Host pathways responsible for the production of AMPs including the *C. elegans* PMK-1 MAPK cascade represent an important conserved mechanism in controlling pathogen proliferation. Mutations in the *C. elegans* p38 ortholog, *pmk-1*^30^ and the MKK3/6 ortholog, *sek-1* lead to greater susceptibility to pathogens including *C. albicans*^32, 33^. Our work demonstrates that removal of host AMPs decreases overall genetic diversity in *C. albicans* (Figures 1 & 2). This suggests that that the large-scale genomic changes generated in hosts with AMPs presents a strategy that enables *C. albicans* to survive this potent defense mechanism and evade future immune response.

Although counterintuitive, LOH increases genetic diversity by unmasking recessive alleles, leading to phenotypic changes^41^. For example, LOH of drug-resistant alleles of *ERG11*, *TAC1* or *MRR1* increases antifungal drug resistance^42–44^. Similarly, aneuploidy offers a short-term solution that organisms use during adaptation^40, 45, 46^ and have been shown to be advantages under certain conditions including the host environment^14–16^. Because we detect large-scale LOH events and aneuploidy in isolates exposed to healthy hosts, this is suggestive that these genetic changes could represent a mechanism for adaptation to the host.

In addition to the host environment increasing genetic diversity through LOH events, physiological stressors including ROS induce LOH and whole changes in ploidy^25, 28^. Our work also demonstrated higher levels of genome instability in *in vitro* ROS (Fig. 4A). In addition, we found that *C. albicans* associated with *bli-3* hosts unable to produce ROS, had lower LOH frequencies (Figure 1), and fewer genome-wide changes compared to wildtype hosts (Figure 2). Further, we also found that *C. albicans* had less INDELs following *bli-3* host exposure (Figure 3B). These novel results suggest that the *bli-3* host defense pathway generates genome instability and genetic variation in *C. albicans*. Recent work in *E. coli*, demonstrated that when exposed to low levels of ROS, *E. coli* exhibits a priming response in which evolution in ROS occurs faster and cells develop greater resistance as opposed to non-primed cells^47^. Our results suggest that host-produced ROS is priming *C. albicans* allowing for tolerance of greater stress.

*C. albicans* has several mechanisms for detoxify ROS, one of which is regulated by the transcription factor Cap1p. *Cap1Δ/Δ* mutants are more susceptible to *in vitro* ROS killing^48, 49^. However, no study has demonstrated this the impact of the Cap1p transcription factor for detoxifying ROS *in vivo*. Here we show that *cap1Δ/Δ C. albicans* genome instability is increased compared to wildtype *C. albicans* in hosts capable of producing ROS (N2 and *sek-1*) but not in *bli-3* hosts incapable of producing ROS (Fig. 4B). Which suggests that host-produced ROS through the bli-3 dual-oxidase induces genome instability in *C. albicans* and that Cap1p is important for detoxification of host-produced ROS.

Exogenous antioxidants provide an alternative to the cellular mechanisms for detoxifying ROS. Adding antioxidants to Pre-SPO media increased cell viability and decreases genome instability in tetraploid *C. albicans*^25^ and prevented the oxidative stress dependent activation of Yap1p in *Saccharomyces cerevisiae* by lowering the overall redox potential^50^. To address if antioxidants reduced genome instability in our system, we exposed *C. albicans* to the well-studied antioxidant lipoic acid^37^ in healthy hosts and immunocompromised hosts. We found that lipoic acid reduced host-associated *C. albicans* genome instability in healthy and *sek-1* hosts, but not *bli-3* hosts (Fig. 4C). We propose that antioxidants reduce *C. albicans* genome instability only when they are exposed to host-produced ROS.

Our work identifies AMPs and ROS as important conserved innate immune responses that generate genome instability in the fungal pathogen, *C. albicans*. We propose that the generation of genetic variation in response to host-produce ROS and AMPs represents a way in which *C. albicans* can quickly respond to host stressors thus further tolerating these stressors or avoiding further immune attack. However, further studies are necessary to define the impact of host-generated genomic changes on the relationship between the host and the pathogen. Our work has important implications for opportunistic pathogens that are often commensal in healthy hosts but more frequently cause infection in immunocompromised hosts.

## Methods

### Strains and Maintenance

*C. albicans* and *C. elegans* strains for this study are listed in Table 1. Yeast strains were stored at −80°C and maintained on YPD (yeast peptone dextrose; 1% yeast extract, 2% bactopeptone, 2% glucose, 0.004% adenine, 0.008% uridine) at 30°C. Strains were initially struck onto YPD agar plates from frozen glycerol stocks and incubated at 30°C for 48 hours and single colonies used as the “parental strain” in subsequent in vivo experiments. Nematode populations were maintained on plates containing nematode growth media (NGM) with *E. coli* (OP50) for a food source. *C. elegans* were transferred to a new plate containing freshly seeded *E. coli* every three to four days. For genome stability assays, treatment plates were seeded with both *C. albicans* and *E. coli* and supplemented with 0.2 g/L streptomycin to inhibit overgrowth of *E. coli*. For fecundity and genome stability assays, NGM was supplemented with 0.08g/L of uridine and 0.08g/L of histidine to facilitate growth of auxotrophic *C. albicans* strains.

### Host-associated C. albicans genome stability

#### Host preparation

NGM plates are seeded with a mixture of *E. coli* and *C. albicans* 24 h prior to host preparation. To seed plates, single colonies of *C. albicans* were inoculated into 3 mL YPD and incubated overnight at 30°C. Cultures were adjusted with ddH_2_O to a final concentration of 3.0 OD_600_ per mL. Simultaneously, a single colony of *E. coli* was inoculated into 50 mL LB and incubated for 24-48 h at 30°C. The *E. coli* culture was pelleted and washed twice with 1 mL ddH_2_O. The pellet was weighed and diluted to final concentration of 200 mg/mL. For *in vitro* treatments, 250 μL *C. albicans* was spread onto NGM + streptomycin agar plates and incubated overnight at 30°C. For *in vivo* treatment plates, 6.25 μL *C. albicans*, 31.25 μL *E. coli*, and brought to a final volume of 250 μL with ddH_2_O, was spread onto NGM + streptomycin agar plates and incubated overnight at 30°C.

To synchronize *C. elegans* populations, nematodes and eggs were washed off plates with M9 buffer, transferred to 15 mL conical tubes, and pelleted via centrifugation (2 min at 1200 rpm). The pellet was resuspended in 2 mL of 25% bleach, mixed via inversion for 2 minutes, and centrifuged for 2 minutes at 1200 rpm. The pellet was washed twice with 3 mL ddH_2_O and resuspended with 1 mL ddH_2_O. To determine the concentration of eggs, 10 μL was pipetted onto a concave slide, eggs were visualized microscopely and counted, and the suspension was adjusted to a concentration of ~100 eggs per 100 μL with M9.

#### Host-associated yeast extractions

Yeast extractions were performed as described previously ^17^. Hosts exposed to *C. albicans* were washed from plates with 3 mL M9 and pelleted via centrifugation (2 min at 2,000 RPM). The supernatant was removed and the pellet was resuspended with 1 mL 3% bleach, transferred to a microcentrifuge tube, and incubated for three minutes and subsequently centrifuged for 1 min at 12,000 rpm. The supernatant was removed and washed with 1 mL of M9 and centrifuged for one minute at 12,000 rpm. The wash was repeated two more times to ensure all bleach was removed. 100 μl aliquots of nematode suspension were transferred to 0.6 mL clear microtubes for manual disruption with a motorized pestle. After one minute of manual disruption, the worm intestine solution was then diluted accordingly with an M9 buffer and plated on YPD + 0.034mg/L chloramphenicol to select prevent any bacterial colonies from arising.

### GAL1 Loss of Heterozygosity assay

#### In vitro

Single colonies of *C. albicans* were inoculated in 3 mL YPD grown overnight at 30°C and subsequently diluted to 3 OD in ddH_2_O. 250 μL was plated and spread onto NGM + streptomycin plates and incubated overnight at 30°C and transferred to 20°C for four days. On day four, yeast cells were washed off with ddH_2_O, harvested by centrifugation, washed once with ddH_2_O, resuspended in 1 mL of ddH_2_O and serially diluted for single colony growth. To determine the total cell viability, 100 μL of 10^−6^ dilution was plated onto YPD and grown for 48 hours at 30°C. To identify cells that lost *GAL1*, 100 μL of 10^−2^ and 10^−3^ dilution was plated onto 2-deoxygalactose (2-DOG; 0.17% yeast nitrogen base without amino acids 0.5% ammonium sulfate, 0.0004% uridine, 0.0004% histidine, 0.1% 2-deoxygalacose, 3% glycerol) and colony forming units (CFUs) counted following 72 hours incubation at 30°C.

#### In vivo

The approach was very similar as the *in vitro* LOH assay described above, with the following changes. A population of ~100 nematodes were plated on each treatment plate containing both *C. albicans* and *E. coli.* On day four, yeast were extracted as described in the previous section. A dilution of 10^−1^ and 10^−2^ was plated on YPD + chloramphenicol to enumerate total growth and undiluted cells were plated on 2-DOG to select for the loss of *GAL1.* Three technical replicates were used for each *C. albicans* strain for both *in vitro* and *in vivo* experiments. At least three biological replicates were used for each genome stability assay.

### Lipoic Acid Experiments

*α*-Lipoic acid (Sigma-Aldrich #1077-28-7) was dissolved in 100% ethanol and added to NGM media containing 0.2 g/L streptomycin sulfate to a final concentration of 25 μM. LOH assays were performed with *α*-Lipoic acid as described above.

### Hydrogen Peroxide Exposure and Genome Stability

Single colonies of *C. albicans* were inoculated in either 2 ml of YPD or in 2 ml of YPD containing 5 mM H_2_O_2_, grown for 20 h at 30°C. Cultures were centrifuged at 2000 rpm for 2 minutes. The supernatant was removed, and the pellet was washed once with 1 ml of ddH_2_O. Cultures were serially diluted for single colony growth. Loss-of-heterozygosity assays were performed to determine the frequency of LOH. Single colonies were randomly picked from YPD plates and genome size was assessed via flow cytometry.

### Whole genome sequencing and analysis

Genomic DNA was isolated with phenol chloroform as described previously^53^. Whole genome sequencing was performed through the Microbial Genome Sequencing Center using a single library preparation method based on the Illumina Nextera kit. Libraries were sequencing using paired end (2 × 150 bp) reads on the NextSeq 550 platform. Adaptor sequences and low-quality reads were trimmed using Trimmomatic (v0.39 LEADING:3 TRAILING: 3 SLIDINGWINDOW: 4:15 MINLEN: 36 TOPHRED33)^54^. All reads were mapped to the phased *C. albicans* reference genome^55^ using the haplotypo python script ‘mapping.py’. This tool uses the Burrows-Wheeler Aligner MEM (BWA v0.1.19) algorithm to align the sequencing reads to the reference genome followed by Samtools (v0.1.19) to sort, mark duplicates, and create a BAM file. Variant files were created using the haplotypo python script ‘var_calling.py’ using the BCFtools calling method.

Identification of aneuploidy, CNVs, and LOH were conducted using whole genome sequencing data and the Yeast Mapping Analysis Pipeline (YMAP). BAM files were uploaded to YMAP and plotted using the Candida albicans reference genome (A21-s02smo8-r09) with corrections for chromosome end bias and GC content^56^.

Identification of SNPs, INDELs and Ts/Tv: Variant files were analyzed using BCFtools ‘stats’ followed by plot-vcfstats to generate a visual representation of the proportion of each type of variant. PCA plot was made be generating GVCF files using haplotypo with the GATK call. To join all of the variant files, the CombineGVCFs GATK module (v.4.0.12.0)^57^ was used. Principle component analysis (PCA) of the SNPs was performed using SNPRelate^58^ in the R/Bioconductor Package.

### Statistical analysis

Statistical analysis was performed using GraphPad Prism 8 software. Data sets were tested for normality using the D’Agnostino & Pearson omnibus normality test.

## Acknowledgments/Author Contributions

We would like to thank Dr. Levi Morran for critical reading of the manuscript. This research is supported by NSF DEB-1943415 (MAH) and Emory University startup funds (MAH).

ACS and MAH designed the study. ACS conducted all of the experiments. ACS and MAH analyzed the data. ACS and MAH wrote, reviewed and edited the manuscript.

## Supplemental

**Table S1:**
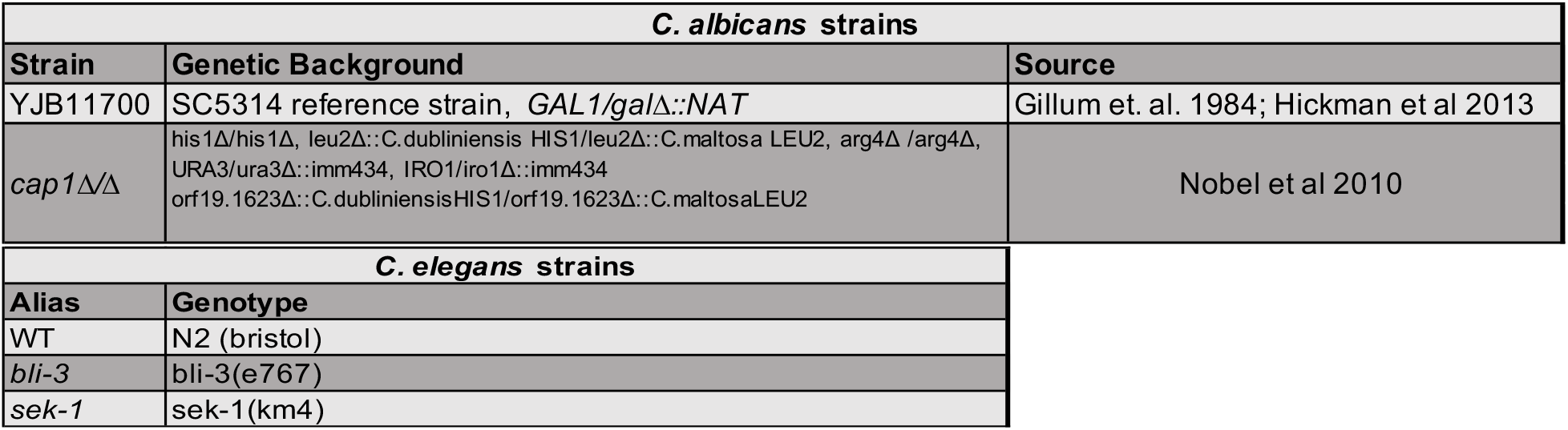
List of strains used in this study

**Figure S1:**
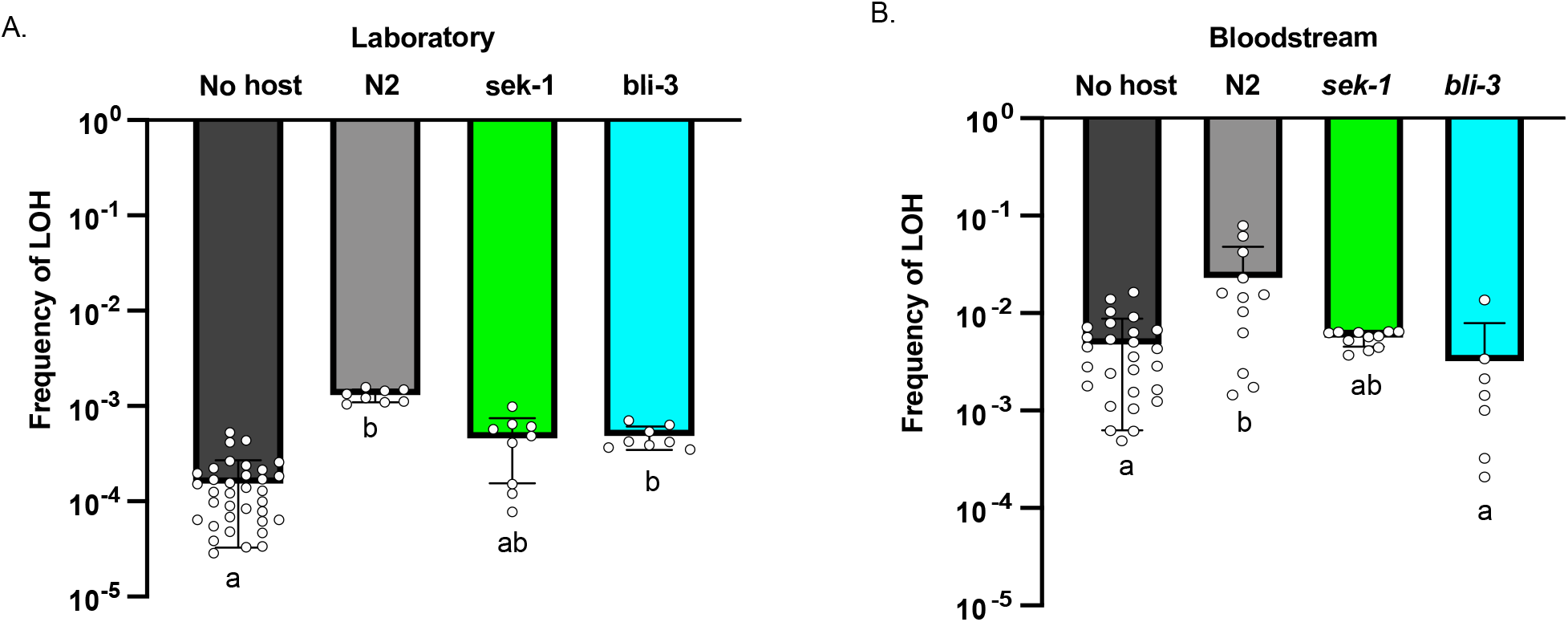
**A)** No host (dark grey, n = 35) and host-associated *GAL1* LOH frequencies for laboratory *C. albicans* extracted from wildtype (N2) (grey, n = 8), *sek-1* (green, n = 9), and *bli-3* hosts (blue, n = 8). Plotted are the mean and standard deviation. Symbols represent individual measurements. **B)** No host (dark grey, n = 27) and host-associated *GAL1* LOH frequencies for a clinical bloodstream strain of *C. albicans* extracted from wildtype (N2) (grey, n = 12), *sek-1* (green, n = 11), and *bli-3* hosts (blue, n = 7). Plotted are the mean and standard deviation. Symbols represent individual measurements. Treatments that share letters are not significantly different, whereas treatments with different letters are statistically different according to a Kruskal-Wallis test with post hoc Dunn’s multiple comparisons (p < 0.05).

**Figure S2:**
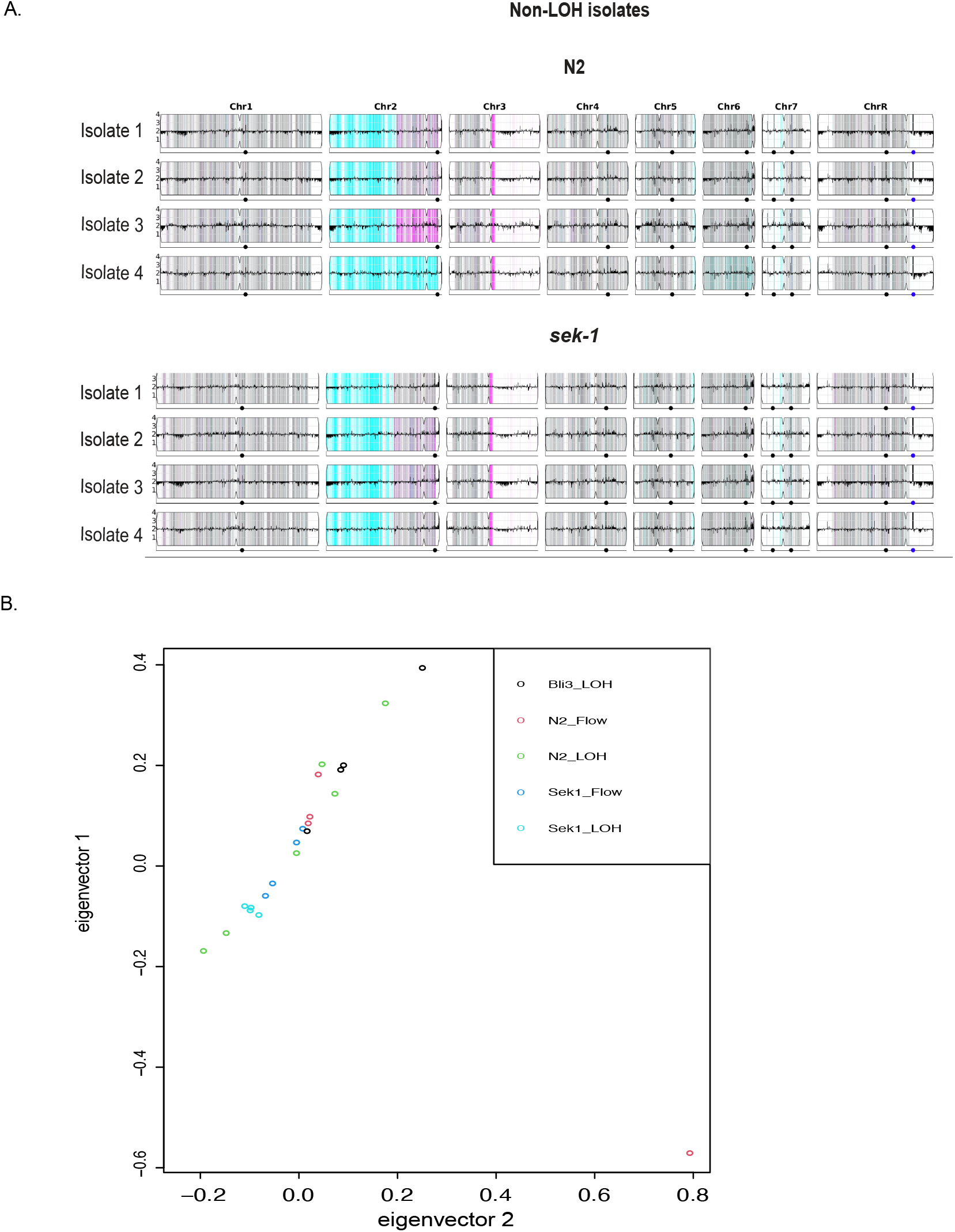
**A)** Following exposure to N2 (wildtype) and *sek-1 C. albicans* was extracted and colonies were chosen at random for DNA extraction and whole-genome sequencing. YMAPs showing the copy number and allelic ratio for each isolate following host exposure. **B)** Eigenvector plot displaying the top two eigenvectors (7.7% and 7.2%)

**Figure S3:**
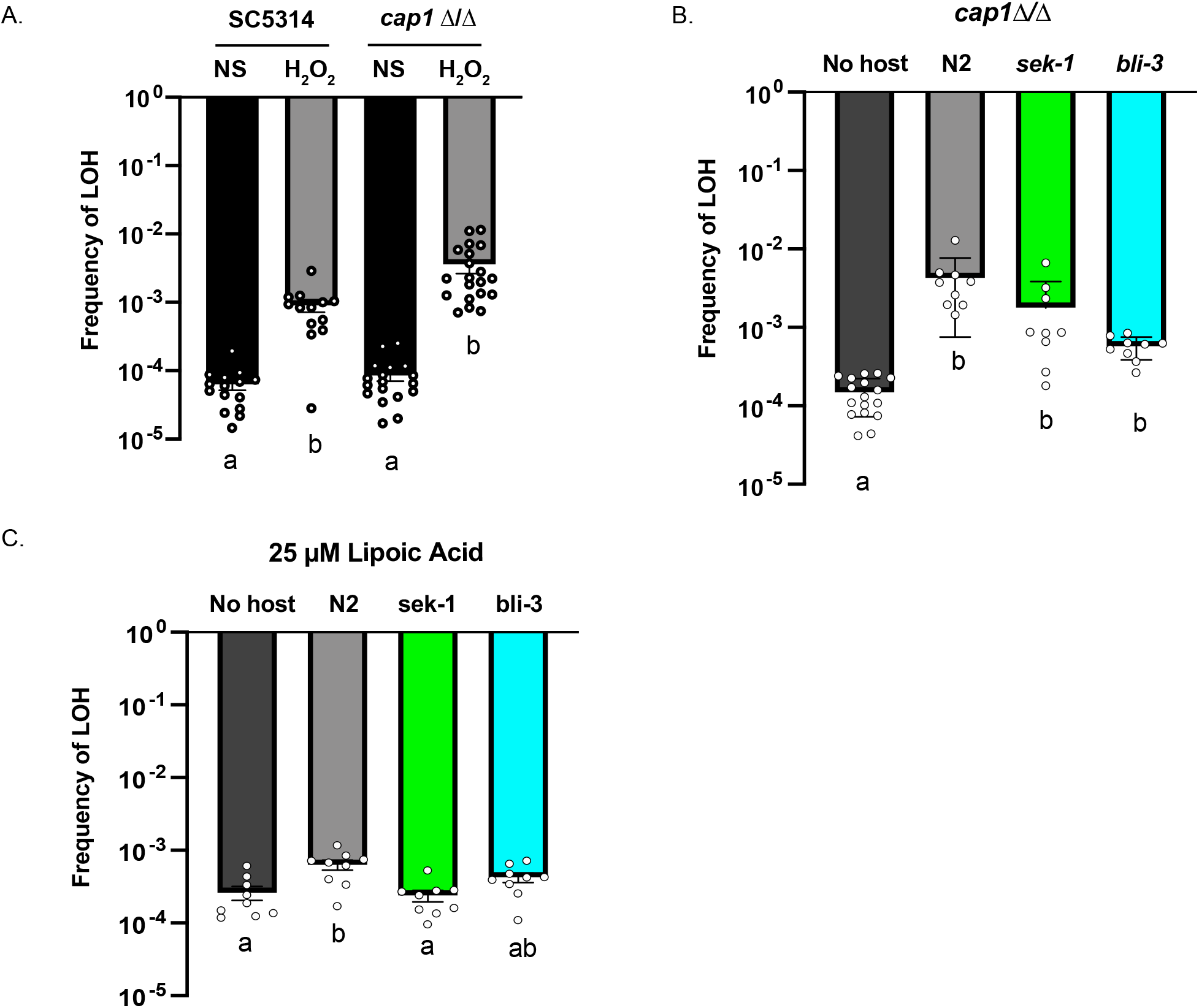
**A)** A) Plotted are the LOH frequencies *C. albicans* exposed to 5mM H_2_O_2_ or no stress for 24 h. Plotted are the means and SD for both SC5314 (no stress: black, n = 15; H_2_O_2_: grey, n = 13) and *cap1Δ/Δ* (no stress: black, n = 18; H_2_O_2_: grey, n = 20). Each data point represents an individual measurement. **B)** No host (dark grey, n = 17) and host-associated *GAL1* LOH frequencies for *C. albicans* extracted from wildtype (N2) (grey, n = 9), sek-1 (green, n = 9), and bli-3 hosts (blue, n = 9). Plotted are the mean and standard deviation. Symbols represent individual measurements. **B)** No host (dark grey, n = 9) and host-associated *GAL1* LOH frequencies for a *C. albicans* extracted from wildtype (N2) (grey, n = 9), sek-1 (green, n = 9), and bli-3 hosts (blue, n = 9) with 25 μM of alpha-lipoic acid. Plotted are the mean and SD. Symbols represent individual measurements. Treatments that share letters are not significantly different, whereas treatments with different letters are statistically different according to a Kruskal-Wallis test with post hoc Dunn’s multiple comparisons (p < 0.05).

